# Linking folding dynamics and function of SAM/SAH riboswitches at the single molecule level

**DOI:** 10.1101/2023.03.08.531827

**Authors:** Ting-Wei Liao, Lin Huang, Timothy J. Wilson, Laura R. Ganser, David M. J. Lilley, Taekjip Ha

**Affiliations:** Department of Biophysics, Johns Hopkins University, Baltimore, MD, USA; RNA Biomedical Institute, Sun Yat-Sen Memorial Hospital, Guangzhou, China; School of Life Sciences, University of Dundee, Dundee, United Kingdom; Howard Hughes Medical Institute, Baltimore, MD, USA

## Abstract

Riboswitches are found in the 5’-UTR of many bacterial mRNAs. They function as cis-acting regulatory elements that control downstream gene expression through ligand-induced conformational changes. Here, we used single-molecule FRET to map the conformational landscape of the SAM/SAH riboswitch and probe how co-transcriptional ligand-induced conformational changes of this translational switch alter ribosome accessibility. The folding of the riboswitch is highly heterogenous, indicating a complex and rugged conformational landscape that enables sampling of the ligand-bound conformation even in the absence of the ligand. Upon ligand binding, the landscape shifts towards the ligand-bound conformation. Mutations at key stabilizing structures alter the ligand-free folding behavior and decrease ligand responsiveness. We also explored translational regulation through folding kinetics by utilizing short oligonucleotides to probe the accessibility of the Shine-Dalgarno sequence within the riboswitch. Additionally, we employed a helicase-based vectorial folding assay to simulate co-transcriptional folding. We find that a competition between ligand binding and ribosome binding is fined tuned via the kinetics of folding. During transcription, the riboswitch takes minutes before reaching equilibrated conformations, and such slow equilibration decreases the effective ligand affinity. Overall, our data demonstrate the significance of conformational polymorphism in RNA function, emphasizing the utilization of complex folding landscapes for regulating ribosome accessibility through ligand induction. Furthermore, we provide direct evidence on how folding kinetics modulate this regulation process.

## Introduction

Riboswitches are regulatory units of RNA that mediate gene expression in response to binding of specific metabolites. They are widely found in bacteria (1–3) but also exist in archaea (4), plants (5) and fungi (6, 7). To date, more than 40 classes of riboswitches have been discovered, and they bind chemically diverse ligands and contribute up to 4% to the bacteria genetic control, especially in gram positive bacteria. Riboswitches are mostly located at the 5’-untranslated regions (5’-UTR), upstream of the regulated genes, and include an aptamer domain capable of binding a particular metabolite with exceptionally high specificity. The riboswitch adopts a specific fold on binding the ligand, leading to up- or down-regulation of the gene either by altering transcription or translation. Since the riboswitch folds and acts as a regulatory unit during transcription, the timing of ligand binding and conformational change is critical, necessitating investigation into its folding kinetics. Substantial research has been devoted to the origins of specificity, correlation of sequence and structure (8, 9), folding kinetics (10), and identification of candidates that can be adapted for drug-delivery (11) and in vivo imaging (12).

*S*-adenosylmethionine (SAM)-binding riboswitches comprise one of the largest classes of riboswitches (1). SAM is synthesized from methionine and ATP by SAM synthetase, encoded by the *metK* gene. SAM is an essential co-substrate of methyltransferases, supplying the methyl group for methyl transfer. Once the methyl group is donated, the resulting *S*-adenosyl-L-homocysteine (SAH) is degraded due to its toxicity (13, 14). To maintain SAM concentration, SAM acts as an inhibitor of MetK synthesis (15–18). This regulation is achieved by the SAM-riboswitches, which bind SAM and acts as negative feedback unit for genes in methionine or cysteine biosynthesis. When SAM concentration goes up, expression of genes in methionine or cysteine biosynthesis is reduced by its upstream 5’-UTR adopting translation-off conformation.

Six sub-classes of SAM riboswitches (SAM-I to SAM-VI) have been identified, classified into three families according to their structural features (17–25). In general, SAM riboswitches exhibit strong discrimination between SAM and SAH by electrostatically interacting with the positive-charged sulfonium cation of the SAM molecule, as previously shown by X-ray crystallography and single-molecule methods (20, 32). By contrast, a SAM/SAH riboswitch does not discriminate between SAM and SAH (33, 34).

The ligand binding interactions of this particular SAM/SAH riboswitch have been previously revealed by NMR and X-ray crystallography (33, 34). Binding of SAM or SAH is accompanied by the formation of three new base pairs that extend the helix of the stem-loop P1, and formation of a pseudoknot helix PK (Fig. S1A). The ligand is bound in the major groove of the extended helix, with the methionyl nitrogen and the adenine moiety hydrogen bonded to a specific cytosine nucleobase. The methionine side chain containing the sulfonium of SAM or the thioether of SAH does not make any direct contact with the RNA, which explains the inability to distinguish between the two ligands. Single-molecule FRET (fluorescence resonance energy transfer) (35) was utilized to compare the binding of SAM and SAH and their kinetic characteristics (34), and no significant differences were observed between the two ligands. Although the ligand-bound state and basic kinetics have been characterized, important features such as ligand-free conformations, binding, folding kinetics, and its role in modulating translation initiation activity are still unknown.

Here, we used single-molecule FRET to investigate the ligand-free and ligand-bound conformations of the SAM/SAH riboswitch and map the energy landscape of folding and altered ribosome accessibility. Folding of the riboswitch is highly heterogeneous, suggesting a rugged conformational landscape that allows for sampling of the ligand-bound conformation even in the absence of ligand. The addition of ligand shifts the landscape, favoring the ligand-bound conformation. Mutation studies showed that the PK helix is crucial for determining ligand-free conformations and their ligand responsiveness. We also investigated the accessibility of the ribosomal binding site under two scenarios: (1) pre-folding of the riboswitch in advance, and (2) vectorial release of the RNA by mimicking co-transcriptional folding. Vectorial folding initially favors a less compact conformation, and it takes minutes before conformational redistribution to that of the pre-folded riboswitch. Such slow equilibration decreases the effective ligand affinity. Overall, our studies offer a deeper understanding of the complexity of the folding process, revealing the mechanism by which the riboswitch adapts its folding pattern in response to ligand and modulate ribosome accessibility and how co-transcriptional folding influences these processes.

## Results

### Heterogeneous folding energy landscape of ligand-free riboswitch

First, we determined the conformational dynamics of the SAM/SAH riboswitch in the absence of ligand. Single riboswitch molecules were tethered to the quartz slide by hybridization of a 3’ extension to an oligonucleotide carrying a Cy5 acceptor at its 3’ terminus. The riboswitch construct has a Cy3 donor attached internally within the loop region such that FRET efficiency, *E*_FRET_, between the two fluorophores can be used to distinguish between conformations. We anticipated two major conformations: the open state with a stem-loop structure previously determined by in-line probing and the closed state with a H-type pseudoknot (Fig. 1A, (29)). The closed conformation likely has a global conformation similar to the crystal structure of the liganded riboswitch that showed the 8-bp extended P1 helix and 5-bp pseudoknot (PK) helix coaxially stacked with each other (34). We previously showed that Cy3 labeling in the loop region does not perturb folding (34).

**Figure 1.**
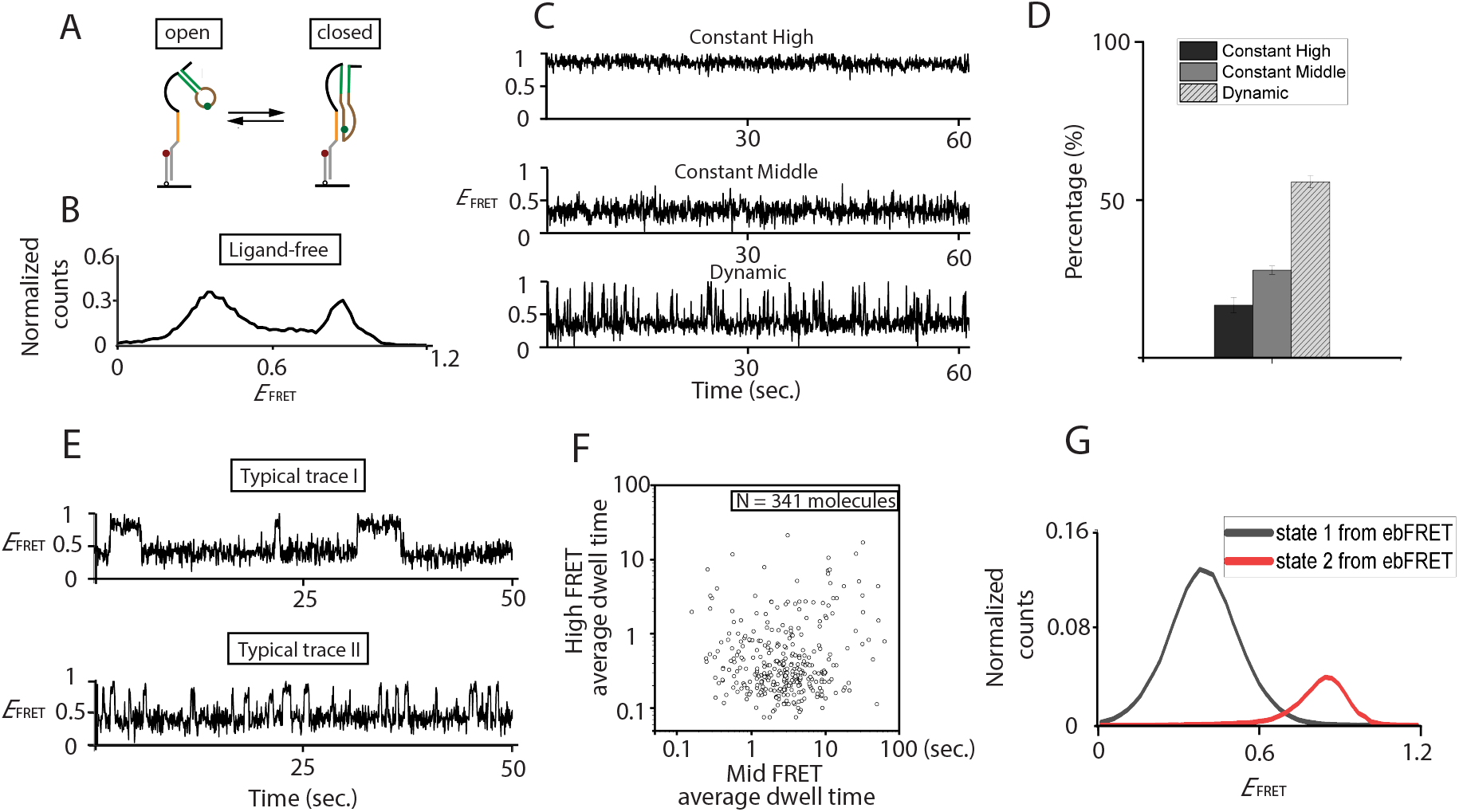
Studies of ligand-free conformations of the SAM/SAH riboswitch by single-molecule FRET. (**A**) A scheme showing the probable folding of SAM/SAH riboswitch RNA. An 18 nt DNA molecule with a 3’ Cy5 acceptor (red circle) was attached via its biotinylated 5’ terminus to a quartz slide. Cy3 donor (green circle) was attached to the bulged nucleotide in the PK helix of the riboswitch, and an 18 nt 3’ DNA extension complementary to the surface-attached DNA allowed the riboswitch to be tethered to the slide. If the pseudoknot helix is not formed the fluorophores should be separated (the open conformation with mid FRET efficiency) whereas in the folded structure the fluorophores should be much closer (the closed conformation with high FRET efficiency). (**B**) Distribution of FRET efficiencies (*E*_FRET_) for SAM/SAH riboswitch molecules corresponding to the open and closed conformations. (**C**) Characteristic traces of *E*_FRET_ as a function of time recorded. Three representative traces are shown, illustrative of constant high FRET (top), constant mid FRET (middle) and dynamic molecules (bottom) undergoing transitions between states of high and middle *E*_FRET_. (**D**) Histograms showing the relative fraction of constant high FRET (black), constant middle FRET (dark gray) and dynamic molecules (light grey with line). (**E**) Among traces showing dynamic transitioning, two characteristic traces are shown, indicating the kinetics of transitioning is diverse and heterogenous. (F, G) Such diverse transitioning kinetics is then fitted into 2 states transitioning by ebFRET. Individual molecule average dwell time is plotted into log-scale scatter plot (**F**), indicating transitioning heterogeneity. And the high FRET and middle FRET probability within the dynamic population is plotted individually (**G**).

Single-molecule histograms of *E*_FRET_ obtained showed two major peaks, likely corresponding to the open conformation (mid-*E*_FRET_ = 0.4, Fig. 1B) and the closed conformation (high-*E*_FRET_ = 0.84, Fig. 1B), suggesting the ligand-bound conformation is adopted even without ligand. Lowering magnesium concentration reduced the high-*E*_FRET_ population but a significant percentage (> 30%) of high-*E*_FRET_ population remained even in the absence of Mg^2+^ (Fig. S1B), suggesting that monovalent cation alone can stabilize the closed conformation.

Single-molecule time traces of *E*_FRET_ displayed three types of behavior: (i) constant mid-FRET (*E*_FRET_ = 0.4) (ii) constant high-FRET (*E*_FRET_ = 0.84) and (iii) dynamic behavior showing transitions between mid- and high-FRET values (Fig. 1C). The majority (55%, Fig. 1D) of traces showed dynamic behavior, further indicating that even in the ligand-free state the closed conformation is sampled. The interconversion kinetics of the dynamic population was quantified by calculating the average dwell times for high and mid- *E*_FRET_ states for each molecule and visualized as a log-scale scatter plot. The average dwell times were widely spread, spanning up to 3 orders of magnitude (Fig. 1E and Fig. 1F), and the open conformation was longer-lived than the closed conformation in most cases (Fig. 1G). The dynamic transitioning was a long-lasting characteristic with no clear population interconversions to or from constant mid/high-FRET states within our experimental window, up to 50 min long (a typical trace shown in Fig. S2A with zoom-in traces in Fig. S2B-D, with intermittent 30 s exposure every 5 min).

### Ligand binding reshapes the folding energy landscape

Next, we measured riboswitch folding in the presence of SAM ligand. The high-FRET state indeed represents the closed conformation because the cognate ligand SAM increased the population of the constant high-FRET species (Fig. 2A). The fraction of molecules in the high-FRET state vs ligand concentration could be fitted using a simple two-state binding isotherm, yielding a dissociation constant (*K*_d_) of 10 μM, similar to those measured in bulk solution using isothermal calorimetry (34). Notably, a significant fraction (38%) of molecules still exhibited dynamic transitioning even when excess ligand was added. The increase in the constant high-FRET species appeared to occur at the expense of the dynamic species while the population of the constant mid-FRET species remained unchanged (Fig. 2B). This suggests that the molecules already in dynamic exchange with the closed conformation were more readily locked into the closed conformation via ligand binding, while the constant mid-FRET population may be trapped in a misfolded state that is not easily rescued by ligand binding.

**Figure 2.**
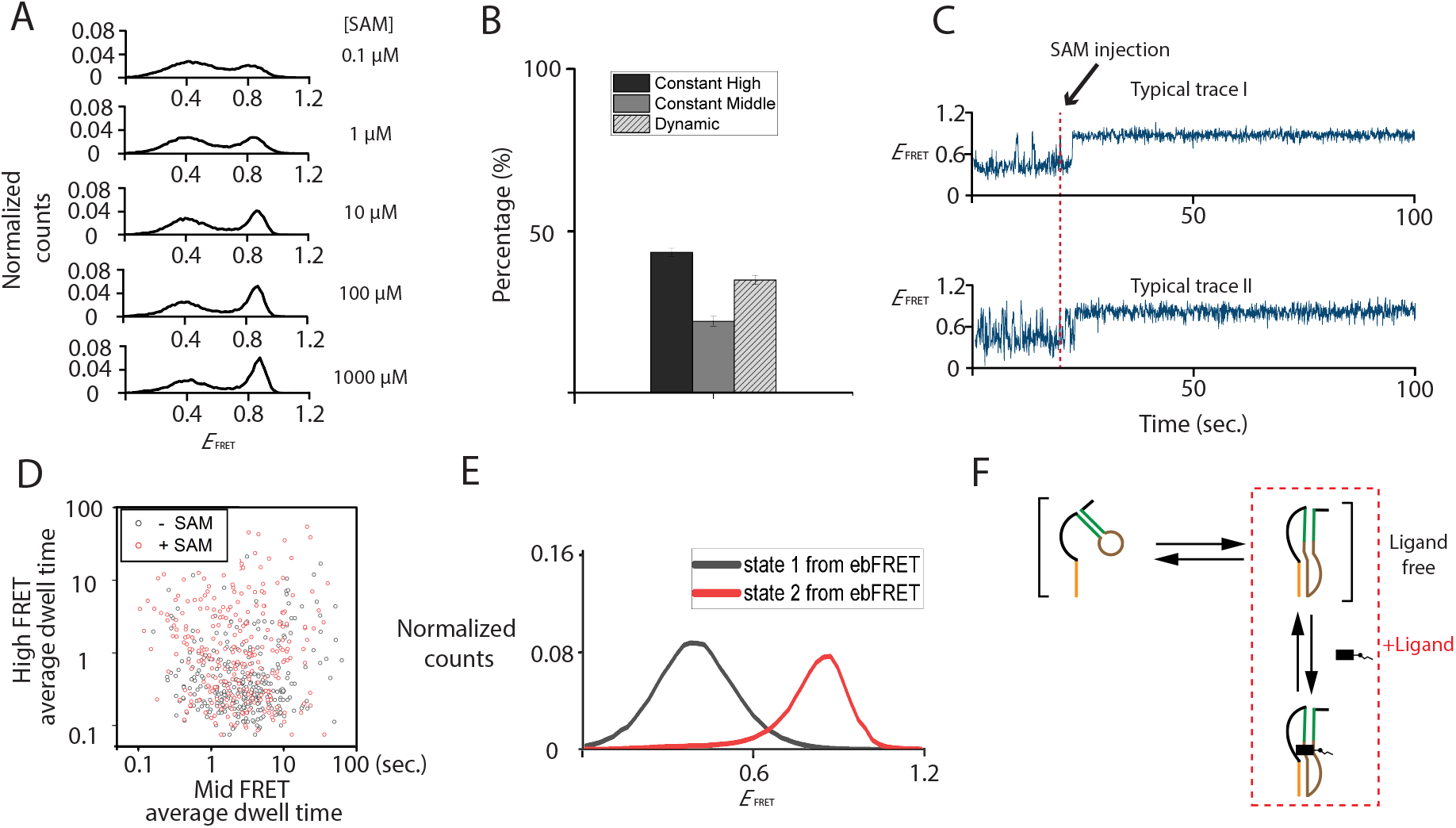
Studies of ligand-induced conformations of the SAM/SAH riboswitch by single-molecule FRET. (**A**) Distribution of FRET efficiencies (*E*_FRET_) as a function of SAM and (**B**) the histograms showing the relative fraction of constant high FRET (black), constant middle FRET (dark gray) and dynamic molecules (light grey with line) in the presence of 1 mM SAM. (**C**) Two typical trajectories of riboswitches showed populations converted from dynamics to constant high FRET while SAM was flowed into the reaction chamber at 20 sec, corresponding to the change of the relative populations in the presence of SAM. (D, E) Among molecules remained transitioning, transitioning kinetics was then fitted into 2 states by ebFRET. Individual molecule average dwell time is plotted into log-scale scatter plot (**D**). And the high FRET and middle FRET probability within the dynamic population is plotted individually (**E**). (**F**) A scheme showing in the presence of ligand, conformations are shifted toward the closed conformations.

To further test our interpretation, we carried out single molecule experiments as the ligand is added via flow. Around 43 % of dynamic species (37 of 86) showed clear locking into the closed conformation after addition of 1 mM SAM (Fig. 2C). We then calculated the average high and mid-FRET state dwell times of each molecule that remained dynamic even after ligand addition. The heterogeneity persisted, but notably the high-FRET dwell time became significantly longer upon ligand addition (Fig. 2D). The heterogeneity increased significantly for the high-FRET dwell time upon ligand addition but remained similar for the mid-FRET dwell time, suggesting that the mid-FRET state is ligand-free (Fig. 2E). A schematic model is presented in Fig. 2F, which illustrates that in the absence of ligand, the riboswitch is in a dynamic equilibrium between folded and unfolded states, and that the addition of a ligand results in a shift towards the closed conformation.

### Structural perturbations provide insights into the folding energy landscape

We next examined alterations in folding behavior caused by mutations that are designed to impact the local structural stability. As shown in Fig. 1A and Fig. 3A, the ligand-bound riboswitch adopts H-type pseudoknot structure with three stabilizing features: (1) P1x: the extension to helix P1, comprising one W-C base pair and two non-W-C pairs, (2) PK: the pseudoknot helix, involving the Shine-Dalgarno sequence, and (3) a triple base interaction (G47:C16-G16) that is part of the PK helix (Fig. 3A). These structural features were abbreviated P1x, PK, and T respectively.

**Figure 3.**
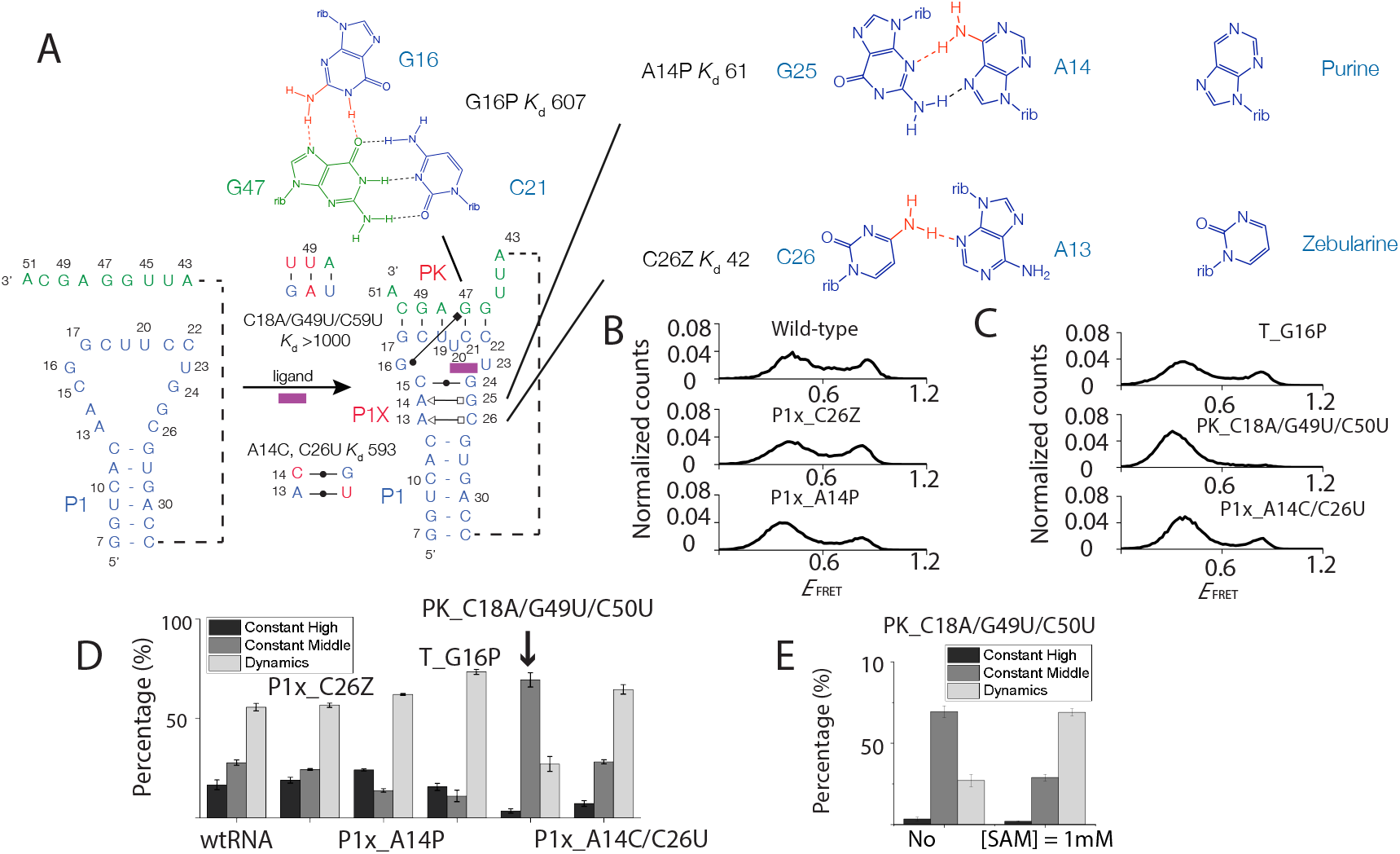
Studies of mutations at local structures affect conformations and ligand responsiveness. (**A**) The ligand-bound riboswitch previously revealed by X-ray crystallography (34) adopts H-type pseudoknot structure with three stabilizing features: (1) P1x: the extension to helix P1, comprising one W-C base pair and two non-W-C pairs, (2) PK: the pseudoknot helix, involving the Shine-Dalgarno sequence, and (3) a triple base interaction (G47:C16-G16). (B, C) Distribution of FRET efficiencies (*E*_FRET_) of the ligand-free conformations of, (**B**) wild-type, P1x_C26Z, and P1x_A14P mutants; (**C**) T_G16P, PK_C18A/G49U/C50U, and P1x_A14C/C26U mutants. All mutations shared similar folding behaviors of constant high FRET, constant middle FRET, and dynamics. (D, E) The histograms showing the relative fraction of constant high FRET (black), constant middle FRET (dark gray) and dynamic molecules (light grey) of wild-type and all mutants (**D**). Among all mutants, PK_C18A/G49U/C50U showed the most different populations both in the absence or in the presence of ligands (**E**).

To perturb the closed conformation, we designed four different mutants, named according to the location of mutation: P1x_C26Z, P1x_A14P, PK_C18A/G49U/C50U, and T_G16P. For P1x mutants, the original base pairing was altered by introducing a modified nucleotide: zebularine (Z: cytosine with N4 removed) or purine (P: adenine with N6 removed) (Fig. 3A). For mutation of the PK helix, two original CG base pairings were replaced with weaker pairings: AU and GU. For mutation of the base triple G16 was replaced by purine, so disrupting the interaction with the Hoogsteen edge of G47 i.e., T_G16P. In choosing the mutation sites, we avoided altering nucleotides that interact directly with the ligand to minimize disruption of the binding site. Additionally, the number of hydrogen bonds removed was kept to a minimum. The positions of these sequence variations can be found in Fig. S3, and the sequences of the mutants are listed in Table S1.

All mutants exhibited an increase in the high FRET population with increasing ligand concentrations, showing that the mutations did not eliminate the ligand’s ability to stimulate riboswitch folding (Fig. S4A-D). The fraction of the high FRET state versus ligand concentration could be fitted using a two-state binding isotherm, yielding *K*_d_ values (Table 1). Mutants that affect the PK helix stability (PK and T mutants) greatly reduced binding affinity: *K*_d_ (PK_C18A/G49U/C50U) > 1 mM; *K*_d_ (T_G16P) = 607 μM. In contrast, mutations at the P1x region had milder effects: *K*_d_ (P1x_C26Z: 41 μM; P1x_A14P: 61 μM).

**Table 1.**
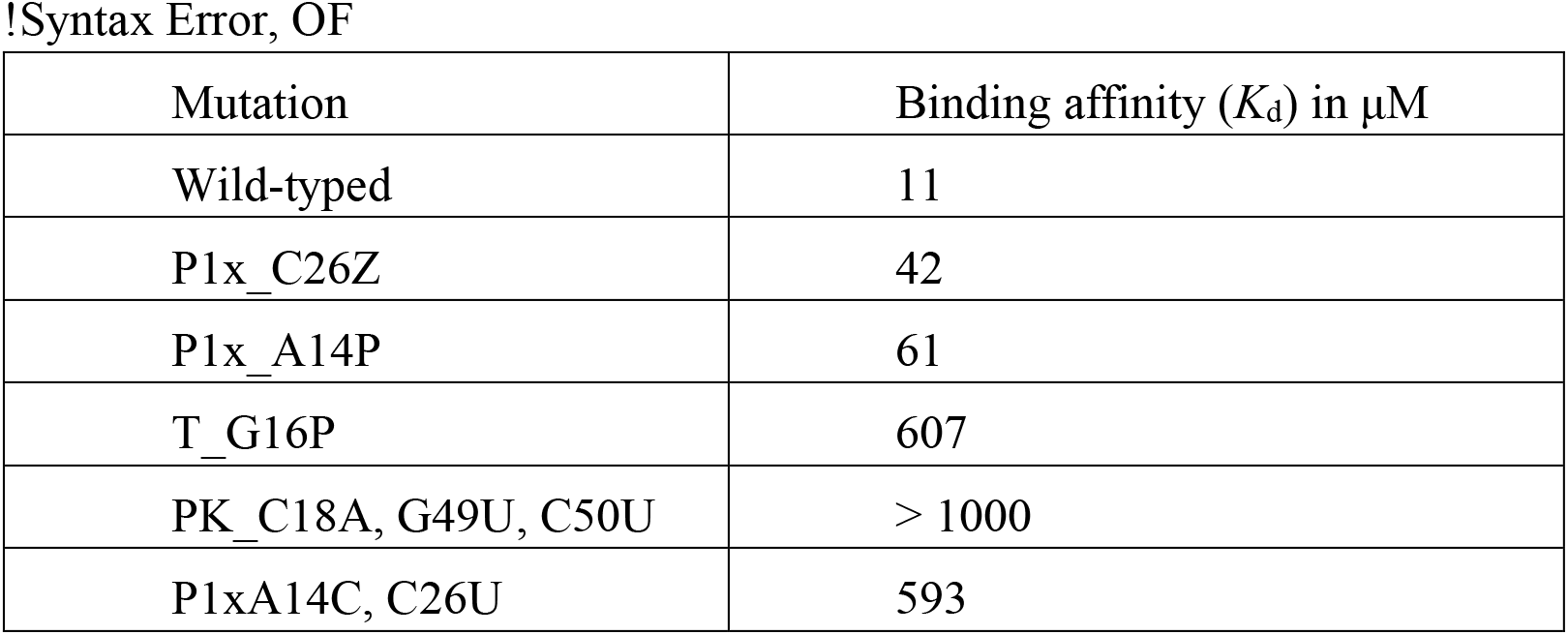
Binding affinity of SAM.

Next, we examined the impact of mutations on ligand-free folding dynamics and ligand responsiveness. In the absence of ligand, all mutants displayed peaks at similar FRET values as the wild type, indicating that open and closed conformations themselves were not significantly altered, but their relative populations changed (Fig.3B and 3C). For example, the destabilization of the closed conformation in the PK_C18A/G49U/C50U mutant leads to a near-complete depletion of the closed conformations (Fig. 3D middle panel). The other three mutants showed a more modest decrease in high FRET peak, by < 3% for P1x_C26Z, 20% for P1x_A14P and 22% for T_G16P (Fig. 3B and 3C). Considering both the binding affinity and the ligand-free conformations, our findings provide evidence for the notion that greater disruptions to the original ligand-free conformations result in greater reduction in binding affinity.

Upon ligand introduction, the general behavior observed in the wild-type riboswitch was observed for all mutants, but the relative populations and their ligand-induced changes were mutation-dependent (Fig. 3D and S5). Notably, mutants that were specifically designed to affect the PK helix stability (PK_C18A/G49U/C50U and T_G16P) exhibited bigger changes in behavior which corresponded well with their greatly reduced binding affinities. As an example, the PK_C18A/G49U/C50U mutant had the constant high FRET population almost depleted, replaced by the dominant constant mid FRET populations (Fig. 3E). Additionally, in the presence of the ligand, the dynamic population became more prevalent at the expense of the constant mid FRET population, while the constant high FRET population still remained nearly depleted (Fig. 3E).

We also tested another mutant P1x_A14C/C26U, this time introducing extra hydrogen bonds to the extended P1 helix (P1x_A14C, C26U) by replacing two original non-WC base pairs (cis sugar-Hoogsteen A13:C26, trans Hoogsteen-sugar A14:G25) with WC base pairs (A13:U26, C14:G25). Despite the introduction of extra hydrogen bonds, the mutants showed a reduction in closed conformation (Fig. 3C bottom panel). In addition, the dynamic species increased in population accompanied by a loss of the constant high FRET species (Fig. 3D). The substitution of the WC base pairs may have disturbed the original stacking geometry of the three extended P1 base pairs, resulting in less stable PK conformation and affecting the base pair C15:G24 that interacts with the ligand, and thus leading to a reduced binding affinity (*K*_d_ = 593 μM). These findings highlight the complex nature of riboswitch folding and how small, localized changes can alter the overall folding equilibrium and responsiveness to ligands, ultimately impacting binding affinity.

### Pseudo-functional readout of ribosome accessibility

We next explored the riboswitch function of blocking translational initiation in a ligand dependent manner by mimicking ribosome binding. The 3’ region of the riboswitch contains Shine-Dalgarno (S-D) site to which the ribosome binds to initiate translation. We used 9 nt oligonucleotides complementary to the S-D region to assess the accessibility of the translation initiation site (Fig. 4A).

**Figure 4.**
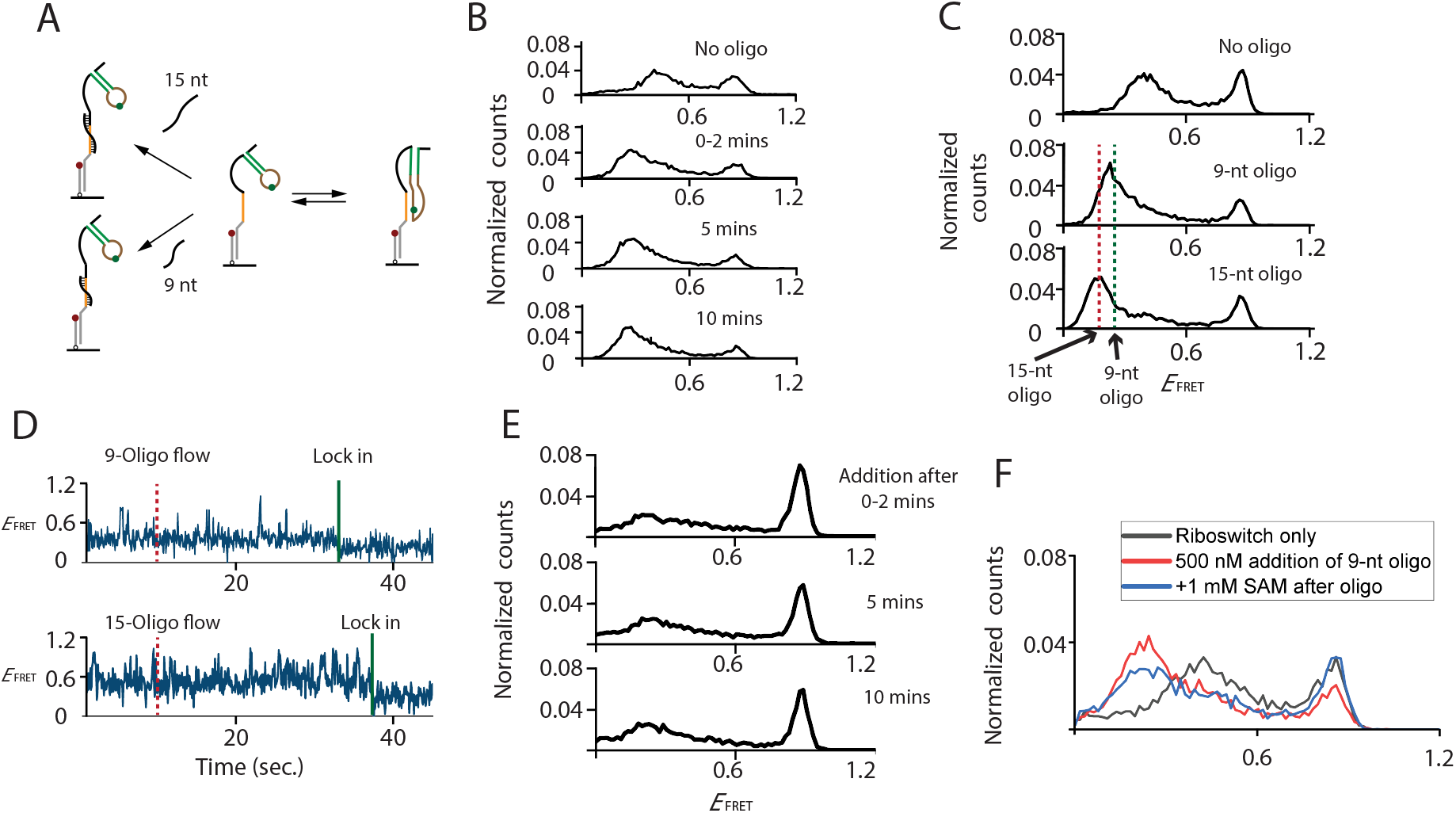
Pseudo-functional studies for assessing the accessibility of the translation initiation site. (**A**) A scheme showing 9 or 15-nt complementary oligonucleotides bind to the open conformation of the SAM/SAH riboswitch. (**B**) Distribution of *E*_FRET_ before and after addition of saturated concentration (= 500 nM) with various time points. (**C**) Distribution of *E*_FRET_ comparing the oligo-free (top), 9-nt (middle), and 15-nt (bottom) oligonucleotide-bound conformations. (**D**) Typical trajectories of riboswitches showed population converted from the mid-FRET to low FRET while oligonucleotides were flowed into the reaction chamber at 12 sec, corresponding to the generated conformations observed in distribution of *E*_FRET_. (**E**) Distribution of *E*_FRET_ after simultaneous addition of 9-nt oligonucleotides (= 500 nM) and SAM (= 1 μM) at various time points, both additions are at saturated concentration. (**F**) Distribution of *E*_FRET_ showing the ligand responsiveness after the riboswitches were pre-bound by oligonucleotides.

In the absence of ligands, adding the oligonucleotides decreased the two major FRET populations (*E*_FRET_ = 0.4 & 0.84) and created a new population with an *E*_FRET_ of approximately 0.22 (Fig. 4B). To identify a condition that is not limited by the rate of oligonucleotides binding to a pre-exposed S-D region, we conducted experiments with varying concentrations of oligonucleotides, and no discernible changes in conformation were observed at or above 500 nM (Fig. S6A). Consequently, all subsequent experiments were performed with the saturating oligonucleotide concentration of 500 nM.

We attribute the state of reduced FRET efficiency to the lengthening of 9 nt S-D region upon riboswitch-oligonucleotides complex formation. A similar experiment using the 15 nt oligonucleotides, mimicking ribosome complex invading 6 nt further into the riboswitch (Fig. 4A), showed the low-FRET peak at a slightly lower value (~ 0.18), consistent with the longer helix that would be formed (Fig. 4C and S6B). Duplex formation was very stable and persisted even after washing away free oligonucleotides (Fig. S6C), in contrast to transient duplex formation observed using a shorter 7 nt long ribosome mimic for 7-aminomethyl-7-deazaguanine (preQ_1_)-sensing riboswitch (36).

We further examined the oligonucleotide binding reaction in real time by flowing in the oligonucleotides during observation. Most of the molecules remained unchanged or showed photobleaching because the binding of the oligo generally took longer than our observation window of ~180 sec. Among molecules showing evidence of oligonucleotide binding, dynamic transitioning was observed before they were locked into the low-FRET state. In addition, most low-FRET states (75% for 9 nt oligo; 91% for 15 nt oligo) were reached from the mid-FRET state (Fig. 4D), likely because the S-D region becomes accessible for oligonucleotide binding only in the open conformation.

### Ligand-induced conformational change outcompetes ribosome mimic binding

We next examined the effect of SAM ligand on the S-D region accessibility and riboswitch folding by simultaneously adding the ligand SAM and oligonucleotide ribosome mimics. Within minutes, the high-FRET population increased at the expense of the mid-FRET population. Only a small fraction of molecules (< 20%) went to the low-FRET conformation corresponding to the ribosome-mimic bound state (Fig. 4E). Therefore, under our experimental condition (1 mM SAM and 500 nM oligonucleotide), ligand binding outcompetes short oligonucleotides binding. Notably, after the addition of ligands and oligonucleotides, the original mid-FRET (0.40) population greatly depleted, suggesting that a subset of mid-FRET species that we found to be ligand-unresponsive is still capable of binding the ribosome mimics.

The riboswitch remained in the high-FRET state for up to 60 min for the 9-nt oligonucleotide. However, for the 15-nt oligonucleotides, ~ 40 % of high-FRET conformation was lost by 10 minutes and by 1 hour, the low-FRET conformation became dominant (Fig. S6D), suggesting that the ligand bound riboswitch can still undergo occasional visits to the open conformation, allowing the 15-nt oligonucleotide to bind to the transiently exposed S-D site. At least for the longer ribosome-mimic, the equilibrium favors the oligo-bound ‘gene-on’ state whereas early in the process, ligand binding can trap the molecule in the closed conformation, momentarily blocking access to the ribosome.

To test if the riboswitch with the ribosome mimic bound to the S-D sequence remains functional, i.e., capable of responding to ligands through conformational changes, we added the ligand after pre-incubation with the oligonucleotides. Whereas ~ 25 % of 9 nt oligo-bound structure were still functional (converted to *E*_FRET_ = 0.84 state), a negligible fraction of the 15 nt bound riboswitch was responsive to the ligand (Fig. 4F and S6E). Therefore, to function as a translational riboswitch, the decision must be made before the arrival of the ribosome, assuming ribosome binding happens only once. In the more realistic case of multiple ribosome molecules arriving and initiating translation in succession, the riboswitch activity can be gradational.

### Mimicking riboswitch folding and ribosome accessibility change during transcription

Because RNA can fold during transcription, riboswitch function should ideally be examined in the context of ongoing transcription (37), and indeed several different methods of inferring co-transcriptional riboswitch folding and function are available (38–43). Here we used the vectorial folding assay where a DNA helicase is used to mimic co-transcriptional RNA folding (38, 43). The riboswitch was hybridized with a complementary DNA oligonucleotide to form an RNA-DNA heteroduplex with a 3’ overhang at the DNA terminus. The same Cy3-Cy5 FRET pair was used to determine the riboswitch conformation (Fig. 5A). A highly processive, engineered DNA helicase, Rep-X (44), is used to unwind the heteroduplex unidirectionally by translocating on the DNA strand in the 3’ to 5’ direction, to release the RNA strand progressively in the 5’ to 3’ direction of transcription and at the speed of transcription, about ~ 60 nt per second (38, 43).

**Figure 5.**
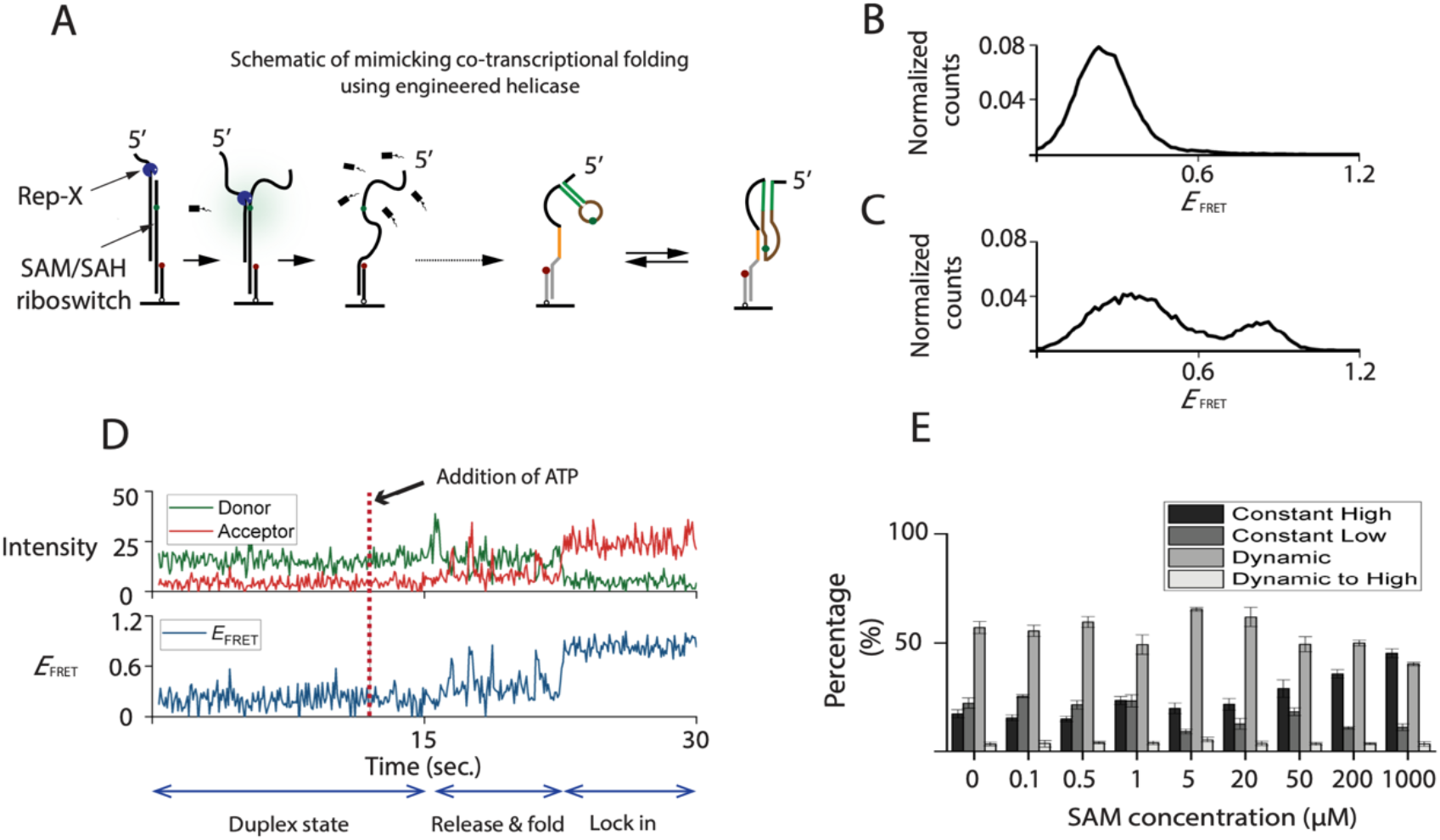
Vectorially folded assays for mimicking co-transcriptional folding. (**A**) A scheme showing the engineered superhelicase Rep-X was preincubated and initiated at designated time for unwinding the heteroduplex. The riboswitch was hybridized with a complementary DNA oligonucleotide to form an RNA-DNA heteroduplex with a 3’ overhang at the DNA terminus. The same Cy3-Cy5 FRET pair was used to determine the riboswitch conformations. (**B**) Distribution of *E*_FRET_ before introducing ATP, corresponding to the heteroduplex conformation. (**C**) Distribution of *E*_FRET_ after vectorially folding, corresponding to the conformations released from the heteroduplex. (**D**) A typical trajectory showing a heteroduplex is unwound and folded into the constant high FRET conformation, where the ATP is flowed in at 12 sec. (**E**) Histograms of relative populations in the presence of various SAM concentrations.

Upon Rep-X addition without ATP, we observed low FRET efficiency (*E*_FRET_ = 0.2) because the fluorophores are separated by the heteroduplex (Fig. 5B). After ATP addition, two new populations centered at 0.4 and 0.84 emerged, corresponding to the open and closed conformations, respectively (Fig. 5C). A representative vectorial folding trace shows two features (Fig. 5D). First, Cy3 intensity showed a transient increase due to protein-induced fluorescence enhancement (45, 46), signifying Rep-X approaching the Cy3 fluorophore on the RNA strand. Second, the heteroduplex was unwound and the riboswitch folding began. We classified the riboswitch folding behavior into four types (Fig. S7A): (I) molecules transitioned from the heteroduplex state to the closed conformation without any detectable intermediate, then remaining there, (II) molecules transitioned from the heteroduplex state to the open conformation, then remaining there, (III) molecules transitioned from the heteroduplex state to one undergoing fluctuations between the open and closed conformations, (IV) molecules transitioned from the heteroduplex to fluctuating states after which they became locked in the closed conformation.

Adding the ligand changed the relative populations of the four types of folding behavior. Type I, direct transition to stable high-FRET state, became more populated at higher ligand concentrations (Fig. 5E) and its fraction vs ligand concentration could be well fitted using a two-state binding isotherm (Fig. S7B), yielding an apparent *K*_d_ value of 108 μM, which is an order of magnitude higher than the *K*_d_ value of 10 μM we observed during ‘refolding’ of preexposed RNA. In addition, the open conformation is more populated at 5 s than at 5 min after ATP addition, most notably in the presence of ligand (Fig. S7C), suggesting that the nascent riboswitch requires more than a few seconds to reach the steady-state conformational distribution, likely reducing its ligand-binding affinity during transcription.

### Vectorial folding tilts the balance toward ribosome mimic binding

In the pseudo-functional read of ribosome accessibility where ligands and ribosome-mimic oligos were introduced to pre-folded RNA, we found that ligand binding is kinetically favored over ribosome mimic binding. To test if this result holds also for vectorial folding where nascent riboswitch may not reach the steady state conformation quickly, we included saturating concentration of 9 nt or 15 nt oligonucleotides during vectorial folding (Fig. 6A). Without ligand, the closed conformation was rarely observed, likely because the S-D region revealed through heteroduplex unwinding is bound by the ribosome mimic before the aptamer can fold into the closed form (top histogram of Fig. 6D and 6E). With ligands, the closed conformation was obtained in a ligand-concentration dependent manner. The efficacy of the ligand in converting the riboswitch to its closed conformation state was diminished in the vectorial folding condition as compared to the refolding condition for both ribosome mimics (Fig. 6B-E). The nascent riboswitch likely first adopts the open conformation, which facilitates ribosome mimic binding, and thus reducing the mutually exclusive ligand bound conformation.

**Figure 6.**
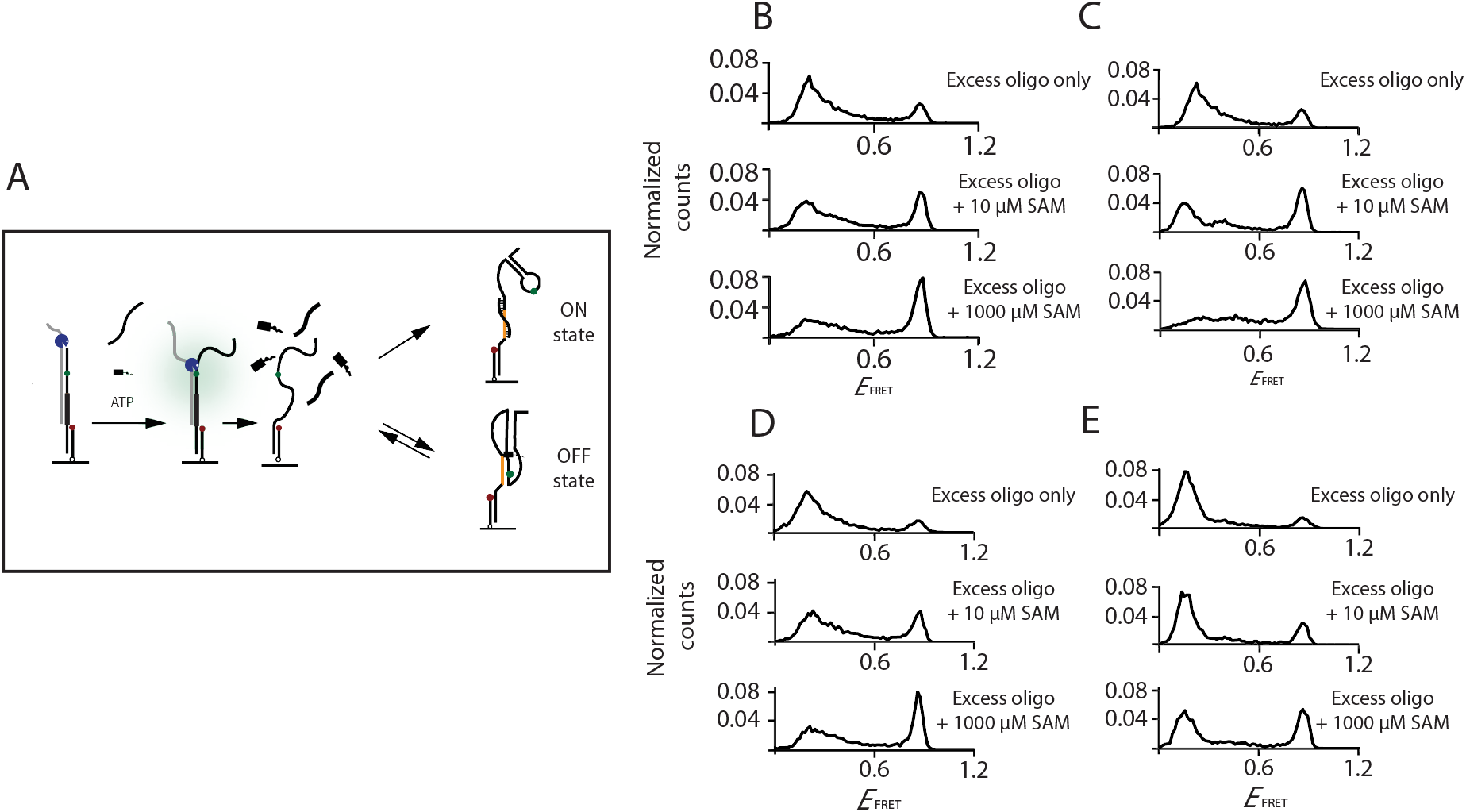
Pseudo-functional studies for assessing the accessibility of the translation initiation site during vectorially folding. (**A**) A scheme showing SAM, oligonucleotides, and ATP are flowed in simultaneously for simulating competition over mutually exclusive conformations during co-transcriptional folding. The oligonucleotide-bound state is termed “ON” state, indicating translation can be initiated, whereas the ligand-bound state is termed “OFF” state, indicating the Shine-Dalgarno site is blocked. (**B**) Distribution of *E*_FRET_ for pre-folded riboswitches under simultaneous addition of 9-nt oligonucleotides and various concentrations of SAM. Similar competitions between oligonucleotides and ligands were carried out while the riboswitches are vectorially folded, and distribution of *E*_FRET_ with various SAM concentrations is shown in (**C**). Similar competition experiments were carried out with longer 15-nt oligonucleotides, for pre-folded competition, (**D**) distribution of *E*_FRET_ with various SAM concentrations; for vectorially folded competition (**E**) distribution of *E*_FRET_ with various SAM concentrations.

## Discussion

We propose a model describing the folding scheme and its energy landscape based on our findings of multiple populations of static folds, open or closed and dynamic switching, and the highly heterogenous switching rates. The FRET values (0.4 and 0.84) of the switching molecules were indifferentiable from those with static conformations, suggesting there are significant structural resemblance. We were surprised that the majority (~ 55%) of ligand-free riboswitches showed dynamic switching between the closed and open conformations. Most of the dynamically switching molecules are responsive to ligand, either by population conversion to the static closed conformation or rate alterations. The observed heterogeneity is likely to be a property inherent to the riboswitch because our constructs with their modifications for surface tethering and fluorescence imaging showed comparable binding affinity to what was determined from unmodified RNA in bulk solution. Furthermore, all five single-site mutants we tested show similarly heterogeneous behavior with only their relative populations and kinetics changed.

Mutants examined in this study showed that not only the populations of static and dynamic populations were strongly affected, but the rates of switching between conformations changed (Figure S8). We speculate that any incomplete base pair formation of the extended P1 stem (E-P1) or PK helix may introduce metastable conformations, leading to heterogeneous folding/unfolding rates for this riboswitch, and potentially for other functional RNAs that also contain the H-type pseudoknot (47–50). Such heterogeneity, if present in vivo, may buffer the riboswitch activity against a wide range of ligand concentrations.

Our findings are most consistent with the previously proposed hybrid model combining conformation selection & induced-fit (10): whereas all conformations are sampled in the absence of ligand (conformation selection), ligand addition repopulates the population ensemble by imparting further stability to the ligand-bound state (induced-fit). A previous SAM-II riboswitch study reported that transient conformational excursions occur in the absence of ligand, suggesting conformational sampling (10). However, they could not determine if those transient conformations were responsive to ligands or how folding and ligand binding are promoted through specific structural motifs.

Relevant to our evaluation of the riboswitch’s accessibility for ribosome mimics, a previous study probed the folding of the 7-aminomethyl-7-deazaguanine-sensing riboswitch using a 7 nt long fluorescently labeled oligonucleotides as translational initiation mimic (46). They observed bursts of probe binding and showed that ligand addition reduces burst duration and extends the intervals between bursts. However, the use of fluorescently labeled probes limited their analysis to sub-*K*_d_ concentrations. By employing unlabeled oligonucleotides, we were able to mimic translation initiation under conditions of saturating ribosome mimic so that the exposure of the binding site is rate-limiting and show that the nascent folds adopted have yet to reach an equilibrium, thus leading to a reduced ligand binding affinity.

In the vectorial folding assay, we observed a decrease in ligand binding affinity, resulting in a reduction in the effectiveness of ligand binding when competing with a ribosome mimic. These differences between pre-folded and vectorially folded riboswitches suggest that the timing of regulatory decision is critical to the effectiveness of the riboswitch and may explain the requirement for higher ligand concentration for effective regulatory control *in vivo* (51). It is possible that there are different modes of regulating accessibility, and the timing of transcription and translation coupling. For tighter regulation, riboswitch needs to reach equilibrium first, thus transcription needs to be carried out in advance of translation. However, when regulation needs not to be tight, the transcription and translation can happen simultaneously.

In conclusion, our studies in this small SAM/SAH riboswitch cast light on the complexity of folding landscape, including individual folding heterogeneity and the dependence of RNA folding kinetics. Our results also have implications on the translational control of the riboswitch; the regulatory decision may be made a nascent riboswitch reaches a folding equilibrium.

## Materials and methods

Riboswitch ligands SAM (A7007), SAH (A9384) were all obtained from Sigma. The RNAs (wide-typed and mutants) are synthesized as described in the following sections. DNA oligonucleotides for mimicking ribosome binding were purchased from Integrated DNA Technologies (Coralville).

### RNA synthesis for single-molecule experiments

The wild-type and mutated SAM/SAH riboswitches for single molecule measurements contains a Cy3 fluorophore attached to the O2’ of U20 generated by Cu^2+^-catalyzed reaction of alkyne-modified RNA with an azide-attached fluorophore (Lumiprobe Corp). The wild-typed RNA had an 18 nt 3’ DNA extension for base-pairing to the anchor DNA, and the complete sequence was (DNA starts with d and underscored): GAUACCUGUCACAACGGCU(U-Cy3)CCUGGCGUGACGAGGUGACCUCAGUGGAG CAA d(<u>ACCGCTGCCGTCGCTCCG</u>), and all the other mutated sequences were showed in Table S1.

The anchor DNA had a 5’-biotin and 3’ Cy5 fluorophore and was complementary to the 18-nt extension of the SAMSAH riboswitch strand. Its sequence was: biotin-CGGAGCGACGGCAGCGGT-Cy5. RNA oligonucleotides were synthesized using t-BDMS phosphoramidite chemistry (52) as described in Wilson et al. (53), implemented on an Applied Biosystems 394 DNA/RNA synthesizer. RNA was synthesized using ribonucleotide phosphoramidites with 2 -Otter-butyldimethyl-silyl (t-BDMS) protection (Link Technologies) (54, 55). Oligonucleotides containing 5-bromocytidine (ChemGenes) were deprotected in a 25% ethanol/ammonia solution for 36 h at 20 °C. All oligoribonucleotides were re-dissolved in 100 μl of anhydrous DMSO and 125 μl triethylamine trihydrofluoride (Sigma-Aldrich) to remove t-BDMS groups, and agitated at 65 °C in the dark for 2.5 h. After cooling on ice for 10 min, the RNA was precipitated with 1 ml of butanol, washed once with 70% ethanol and suspended in double-distilled water. RNA was further purified by gel electrophoresis in polyacrylamide under denaturing conditions in the presence of 7 M urea. The full-length RNA product was visualized by UV shadowing. The band was excised and electroeluted using an Elutrap Electroelution System (GE Healthcare) into 45 mM Tris-borate (pH 8.5), 5 mM EDTA buffer for 12 h. at 150 V at 4°C. The RNA was precipitated with isopropanol, washed once with 70% ethanol and suspended in water or ITC buffer (40 mM HEPES-K (pH 7.0), 100 mM KCl, 10 mM MgCl_2_).

### Ligand titration of pre-folded riboswitches: wild-typed and mutated riboswitches

40 pM of the pre-annealed SAM/SAH riboswitch molecules were immobilized on a neutravidin-functionalized, polymer-passivated surface and free molecules were washed out with T50 buffer. Image buffer containing an oxygen-scavenging system was freshly mixed before measurements, comprising 1 % (w/v) dextrose, 2 mM Trolox, glucose oxidase (1 mg/ml; Sigma-Aldrich), and catalase (500 U/ml; Sigma-Aldrich)] in buffer containing 40 mM HEPES (pH 7.5), 100 mM KCl, 2 mM MgCl_2_. All the ligands were diluted with image buffer immediately prior to measurements. The ligands were incubated for 5 min before imaging. Short movies (duration of 1.5 sec: 20 frames) were collected for 30 field of view for generating distribution of FRET efficiencies (*E*_FRET_). The distribution is then fitted by two individual Gaussian function, and the high *E*_FRET_ ratio is estimated accordingly.

### Single-molecule imaging and data acquisition

Single-molecule FRET data were obtained using a prism-based total internal reflection fluorescence (TIRF) microscope. The Cy3 and Cy5 fluorophores were excited by a 532-nm laser (Coherent Compass 315M) and a 638-nm laser (Cobolt 06-MLD) respectively. The fluorescence emission was collected by a water immersion objective (Olympus NA 1.2, 60×) and recorded by a back-illuminated electron-multiplying charge-coupled device camera (iXON, Andor Technology) with a dual-view setup. The dual-view setup used a long-pass emission filter (Semrock BLP02-561R-25) for eliminating the 532-nm laser, and a notch filter (Chroma ZET633TopNotch) for eliminating the 638-nm laser. The fluorescence emission was separated into donor and acceptor emission by a long-pass dichroic mirror (Semrock FF640-FDi01-25 × 36). The passivated PEG quartz slides and coverslips were purchased from Johns Hopkins Slides Core and were assembled into a reaction chamber. (56) Spots detection, background subtraction, donor leakage and acceptor direct-excitation correction followed our previous protocol (56). Custom codes are available on GitHub (https://github.com/Ha-SingleMoleculeLab).

### Single-molecule data analysis of pre-folded riboswitch

Single-molecule traces showing *E*_FRET_ as a function of time were categorized into three types of behavior. (i) *E*_FRET_ remained middle for the duration of observation, up to 1 min, (ii) *E*_FRET_ remained high for the duration of observation (iii) *E*_FRET_ fluctuated between the middle and high values. To characterize further the dynamic species, the regions of dynamics were collected and analyzed by ebFRET (57) and the two-state dwell time was plotted into log-scale scatter plot.

### Single-molecule data analysis of vectorially folded riboswitch

Single-molecule traces showing the immobilized heteroduplex was unwound were categorized into four types of behavior. We classified the riboswitch folding behavior into four types (Fig. S7A): (I) molecules transitioned from the heteroduplex state to the closed conformation without any detectable intermediate, then remaining there, (II) molecules transitioned from the heteroduplex state to the open conformation, then remaining there, (III) molecules transitioned from the heteroduplex state to one undergoing fluctuations between the open and closed conformations, (IV) molecules transitioned from the heteroduplex to fluctuating states after which they became locked in the closed conformation.

### Pseudo-functional readout of ribosome accessibility of pre-folded riboswitch

40 pM of the pre-annealed SAM/SAH riboswitch molecules were immobilized on a neutravidin-functionalized, polymer-passivated surface and free molecules were washed out with T50 buffer. All the ligand (SAM) and DNA oligonucleotides with a designated concentration was freshly mixed with image buffer containing an oxygen-scavenging system. Short movies (duration of 1.5 sec: 20 frames) were collected for 30 field of view immediately or at designated time (5-min or 1 hr) after injection for generating distribution of FRET efficiencies (*E*_FRET_).

### Sample preparation for the single-molecule FRET measurements

For preparation of the pre-folded assay, 10 μM of the SAM/SAH riboswitch molecule or the riboswitch mutants with internal Cy3 labeled was annealed with 15 μM anchored DNA with Cy5 and biotin label under 1X T50 [10 mM Tris (pH 8.0), 50 mM NaCl) buffer followed by slow cooling from 95°C to room temperature.

For preparing of the vectorial folding assays, 10 μM anchored DNA with Cy5 and biotin label was annealed with 20 μM of the Cy3-labeled SAM/SAH riboswitch and 40 μM complementary DNA oligos (cDNA) with dT30 overhang in 10 μl of 1X T50 [10 mM Tris (pH 8.0), 50 mM NaCl] by incubating the mixture at 95 °C for 1 min, 75 °C for 5 min, 37 °C for 15 min and finally equilibrating at room temperature for 5 min (38, 43).

### Vectorial folding as a mimic of riboswitch folding and ligand binding

Labeled and biotinylated heteroduplexes were immobilized on a neutravidin-functionalized surface and free heteroduplexes were washed out. 50 nM of Rep-X (44) was incubated for 2 min with heteroduplexes in the imaging buffer (40 mM HEPES (pH 7.5), 100 mM KCl, 2 mM MgCl_2_) containing an oxygen-scavenging system, and images (duration of 1.5 sec: 20 frames) were collected for 30 field of view for confirming the heteroduplex conformation. Unwinding was initiated by mixing unwinding buffer with/without ligands at designated concentration. Unless specified otherwise, the unwinding buffer contained 40 mM HEPES (pH 7.5), 100 mM KCl, 2 mM MgCl_2_, 2mM ATP with an oxygen-scavenging system. Buffer with ATP then triggered the pre-bound RepX into unwinding the anchored heteroduplex.

For the real-time observation (“flow-in” experiments) of riboswitch released from heteroduplex, imaging was started 12 seconds before the addition of the unwinding buffer. For characterization of VFA products after helicase unwinding, images were taken after the addition of the unwinding buffer at designated time. The loading and unwinding buffers used during imaging contained additional 1 % (w/v) dextrose, 2 mM Trolox, glucose oxidase (1 mg/ml; Sigma-Aldrich), and catalase (500 U/ml; Sigma-Aldrich).

### Vectorial folding with ligand and ribosome mimic addition simultaneously

Labeled and biotinylated heteroduplexes were incubated with 50 nM Rep-X as previously described, images were taken before addition to confirm the heteroduplex conformations. Ligands, ribosome mimics (oligonucleotides with 9-nt or 15-nt complementary to ribosome binding site), and additional Rep-X (50 nM for 9-nt; 100 nM for 15-nt) were mixed with unwinding buffer. The additional Rep-X is added in order to reduce the competition of free oligonucleotides to the Rep-X pre-incubated before unwinding mixture. This optimized condition is tested with a negative control, where additional Rep-X with dT9 or dT15 were added simultaneously into the heteroduplex, no observable loss of unwinding efficiency in this condition. The negative control experiments were shown in Fig. S10.

