## Supplementary materials for "Linking folding dynamics and function of SAM/SAH riboswitches at the single molecule level"

SUPPLEMENTARY FIGURES
SUPPLEMENTARY TABLES

FIGURE S1

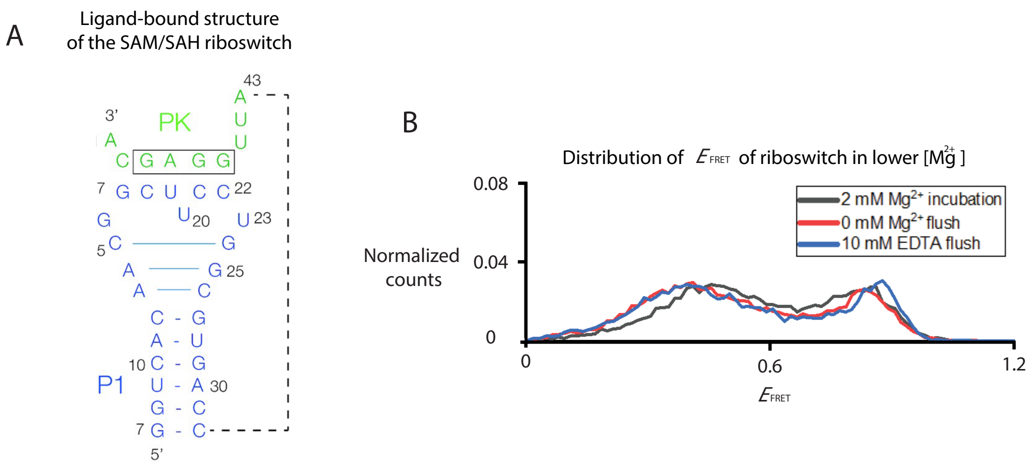


**Figure S1. The secondary structure of the SAM/SAH riboswitch and conformations analysis under depleted magnesium.**

(**A**) The 5′ end of the molecule (blue) adopts a stem-loop structure (P1). The 3′ end (green) base pairs with the center of the loop to generate pseudoknot (PK) helix with five Watson–Crick base pairs, and thus forming an H-type pseudoknot. Three additional base pairs form at the top of the P1 helix. The Shine–Dalgarno ribosome binding site (boxed) is an integral part of the PK helix. We have followed the numbering scheme used by Weinberg *et al.* (33). (**B**) Analysis of conformations under various buffer conditions. Distribution of $E_{FRET}$ incubating in 2 mM $Mg^{2+}$ imaging buffer (black line), after buffer wash with no $Mg^{2+}$ buffer (red line), and additional EDTA incubation to ensure no $Mg^{2+}$ are left in the reaction chamber. Other ingredients of imaging buffer can be found in materials and methods.

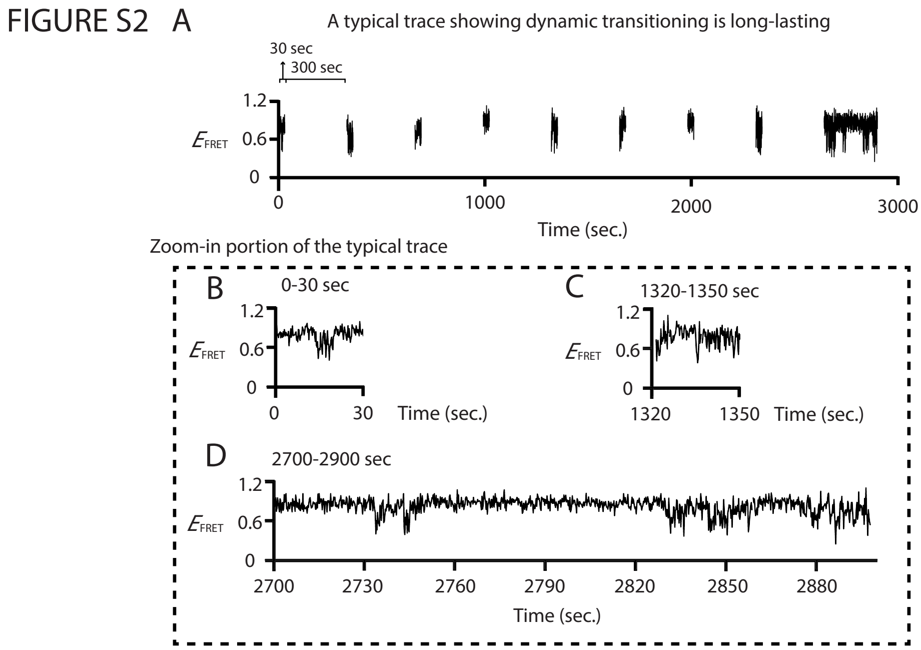


**Figure S2. A typical trajectory of a SAM/SAH riboswitch in the dynamic population.** In order to minimize photobleaching and observe long-lasting transitioning kinetics, molecules are under intermittent 30 s exposure every 5 min except for the last exposure, the total length of the trajectory is around 50 min. (**A**) An overall trajectory showing total 9 sessions of exposure. (B, C, D) Zoom-in sections of the trajectory at 0-30 sec (**B**), 1320-1350 sec (**C**), and the last exposure 2700-2900 sec (**D**).

FIGURE S3

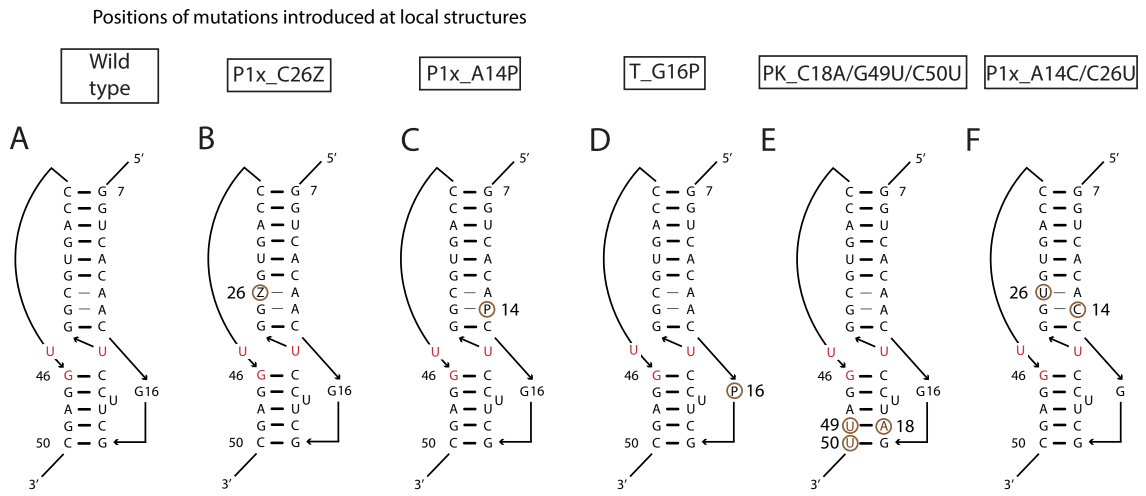


**Figure S3. Simplified schemes of mutations at the local structures.**

This scheme shows mutations introduced at the local structures, emphasizing on the base pairs of the closed conformation. Watson-Crick basepairs are shown in thick horizontal lines, non-WC basepairs are shown in thin horizontal lines. Ligand-interacting nucleotides are shown in red. And individual mutation position is labeled with a number and a brown circle. The sequences neglected in the scheme are all identical for all mutants, full sequences are in Table S1. Individual riboswitch scheme is shown in the following order: wildtype (**A**), P1x_C26Z (**B**), P1x_A14P (**C**), T_G16P (**D**), PK_C18A/G49U/C50U (**E**), and P1x_A14C/C26U (**F**).

FIGURE S4

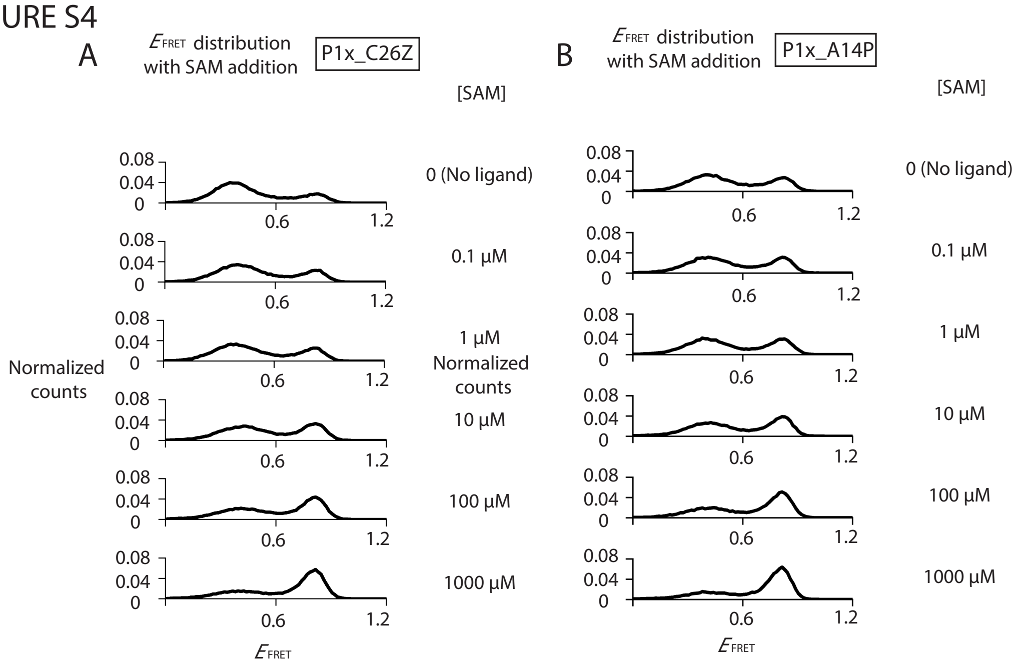


FIGURE S4

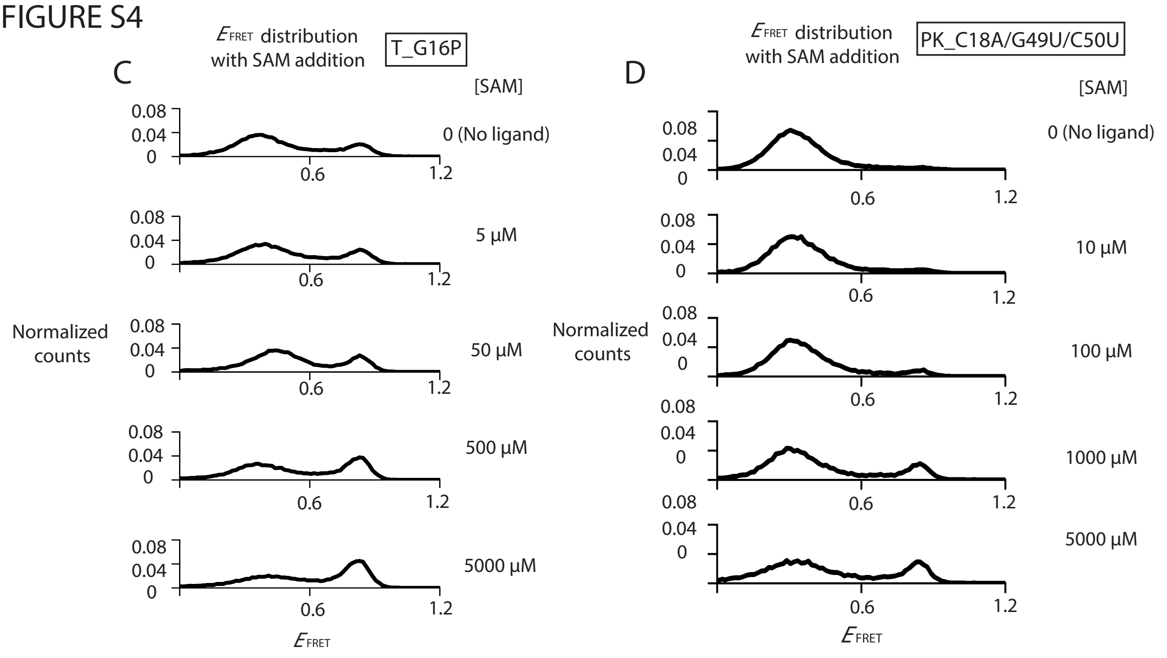

FIGURE S4

E

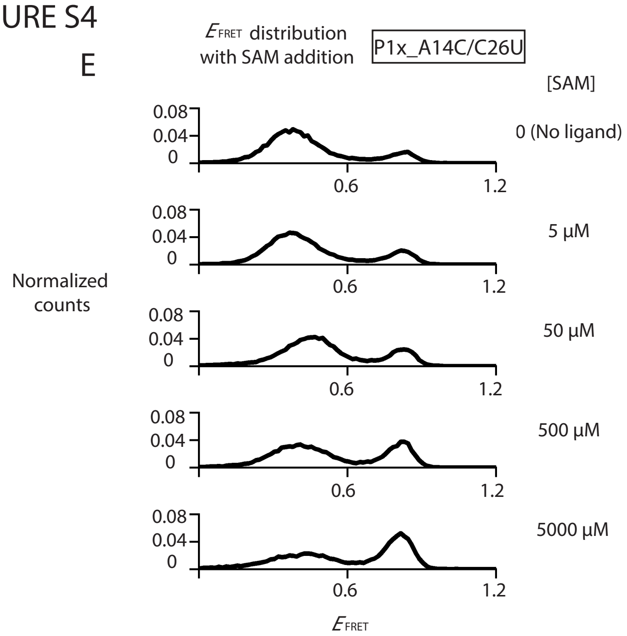


**Figure S4. SAM responsiveness analyses of ligand titration across mutants.**
Individual titration of each mutation showing distribution of $E_{FRET}$ in the context of various SAM concentrations is shown in the following order: P1x_C26Z (**A**), P1x_A14P (**B**), T_G16P (**C**), PK_C18A/G49U/C50U (**D**), and P1x_A14C/C26U (**E**).

FIGURE S5

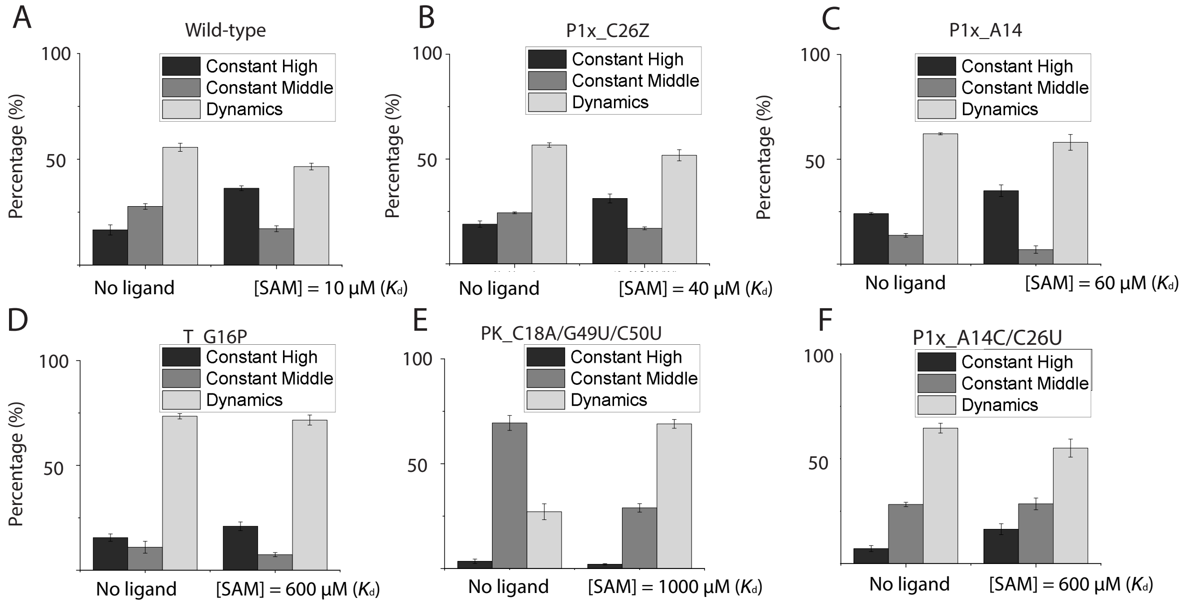


**Figure S5. SAM responsiveness analyses of relative populations across mutants.** Relative populations of individual mutations in the ligand-free and at concentration equal to its individual $K_d$ are shown in the following order: wild-type (**A**), P1x_C26Z (**B**), P1x_A14P (**C**), T_G16P (**D**), PK_C18A/G49U/C50U (**E**), and P1x_A14C/C26U (**F**).

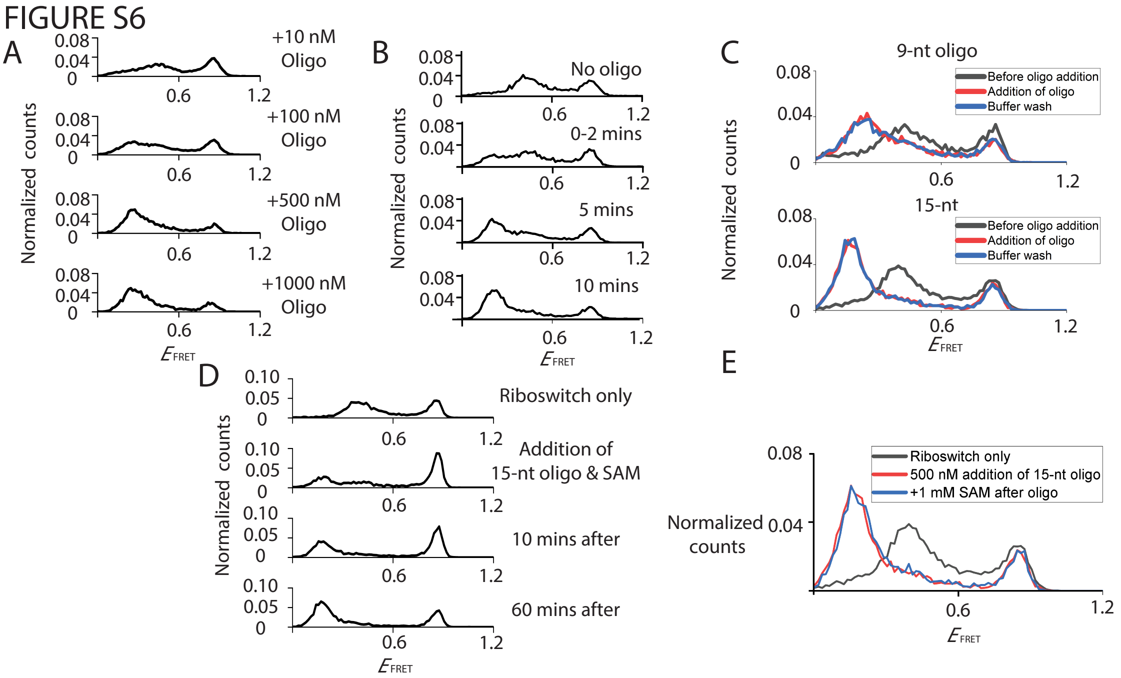


**Figure S6. Pseudo-functional readout for assessing the accessibility of the translation initiation site.**

(**A**) Analyses of various concentrations of oligonucleotides addition. Distribution of $E_{\text{FRET}}$ of titration of 9 nt oligonucleotides at various concentrations, up to 1000 nM. (**B**) Analyses of binding of oligonucleotides. Distribution of $E_{\text{FRET}}$ before and after addition of 15 nt oligonucleotides at saturated concentration (= 500 nM) with various time points. (**C**) Analyses of stability of oligo-bound conformations. Distribution of $E_{\text{FRET}}$ before addition of oligonucleotides (black line), after addition of oligonucleotides (red line), and after wash with oligo-free buffer (blue line). Top panel: 9-nt oligonucleotide, and bottom panel: 15-nt oligonucleotide. (**D**) Analyses of competitions between oligonucleotides and ligands. Distribution of $E_{\text{FRET}}$ after simultaneous addition of 15-nt oligonucleotides and SAM both at saturated concentrations with various time points. (**E**) Analysis of ligand responsiveness of the oligo-riboswitch complex. Distribution of $E_{\text{FRET}}$ before addition of oligonucleotides (black line), after addition of oligonucleotides (red line), and after addition of SAM at saturated concentration (= 1mM).

FIGURE S7

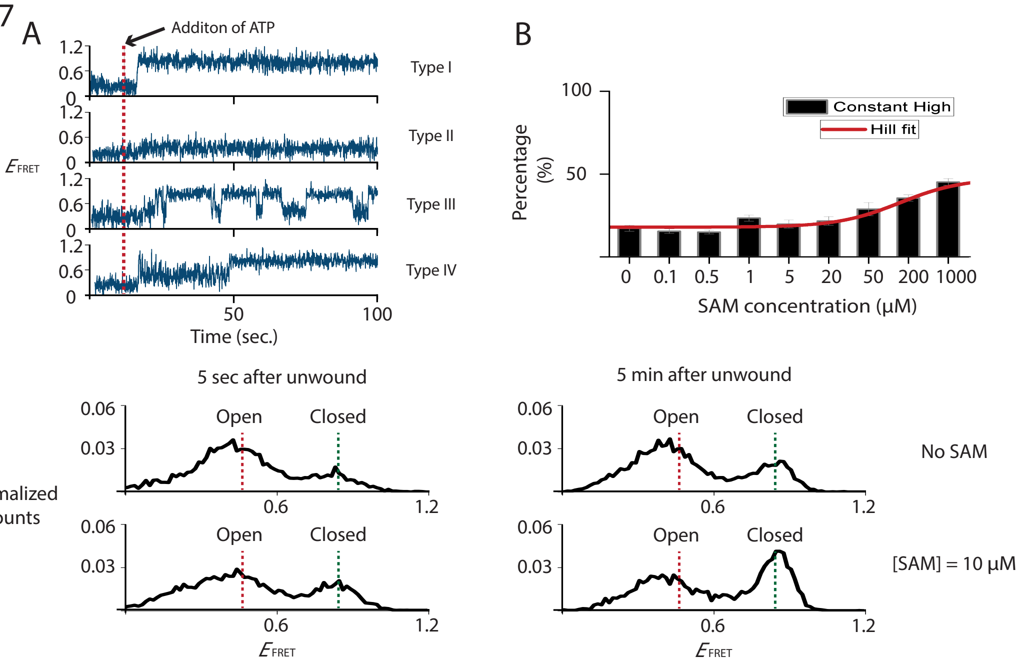


**Figure S7. Folding behaviors of the vectorially folded riboswitch.**

(**A)** Typical trajectories of folding behaviors when ATP is flowed at 12 sec, 4 types of behaviors are observed: (I) molecules transitioned from the heteroduplex state to the closed conformation (II) molecules transitioned from the heteroduplex state to the open conformation (III) molecules transitioned from the heteroduplex state to one undergoing fluctuations between the open and closed conformations, (IV) molecules transitioned from the heteroduplex to fluctuating states after which they became locked in the closed conformation. (**B**) Among relative populations, the fraction of type I, direct transition to stable high-FRET state, vs ligand concentration could be well fitted using a two-state binding isotherm (fitted curve shown in red line). (**C**) Analyses of conformations before and after reaching folding equilibrium under no (top panel) or 10 µM of SAM (bottom panel). Distribution of $E_{FRET}$ after vectorially folding at two different time points: 5 secs after vectorial folding (the left panel), and 5 mins after vectorial folding (the right panel).

FIGURE S8

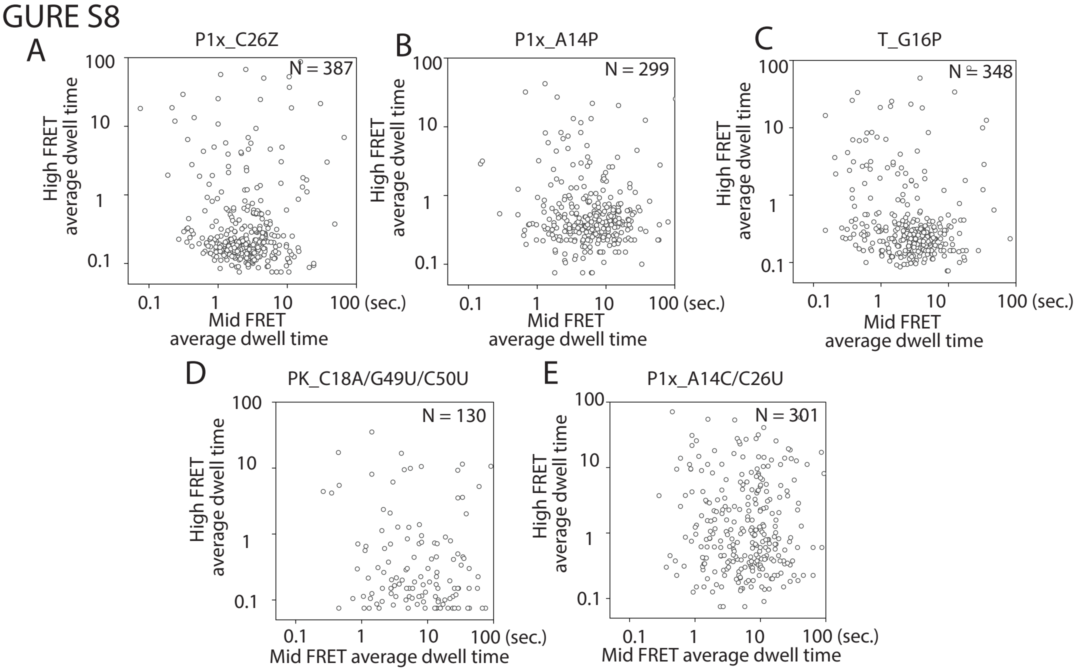


**Figure S8. Analyses of transitioning kinetics among the dynamic population of mutations introduced at local structures.**

Individual molecule average dwell time of the dynamic population is plotted into log-scale scatter plot in the following order: P1x_C26Z (**A**), P1x_A14P (**B**), T_G16P (**C**), PK_C18A/G49U/C50U (**D**), and P1x_A14C/C26U (**E**).

FIGURE S9

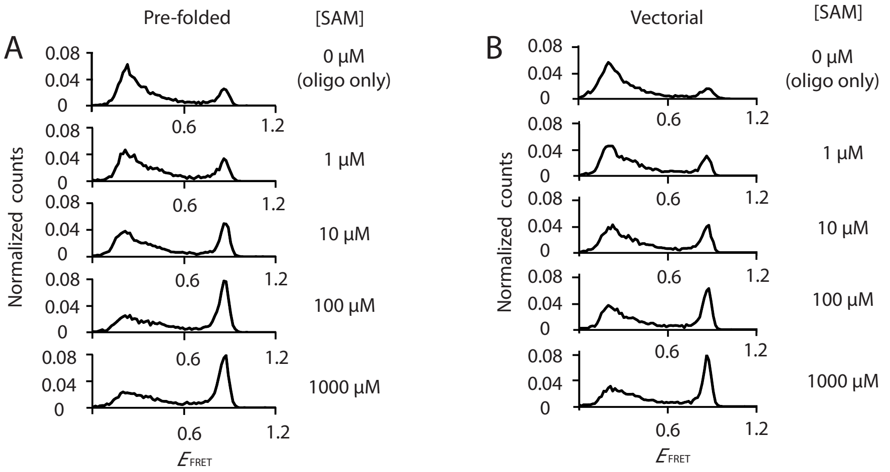


**Figure S9. Pseudo-functional studies for assessing the accessibility of the translation initiation site during vectorially folding.**

Analyses of the ligand responsiveness of simultaneous addition of oligonucleotides and ligands of pre-folded and vectorially folded riboswitches. (**A**) Distribution of $E_{\mathrm{FRET}}$ for pre-folded riboswitches under simultaneous addition of 9-nt oligonucleotides and various concentrations of SAM: 1, 10, 100, 1000 µM. Similar competitions between oligonucleotides and ligands were carried out while the riboswitches are vectorially folded, and distribution of $E_{\mathrm{FRET}}$ with various SAM concentrations: 1, 10, 100, 1000 µM is shown in (**B**).

FIGURE S10

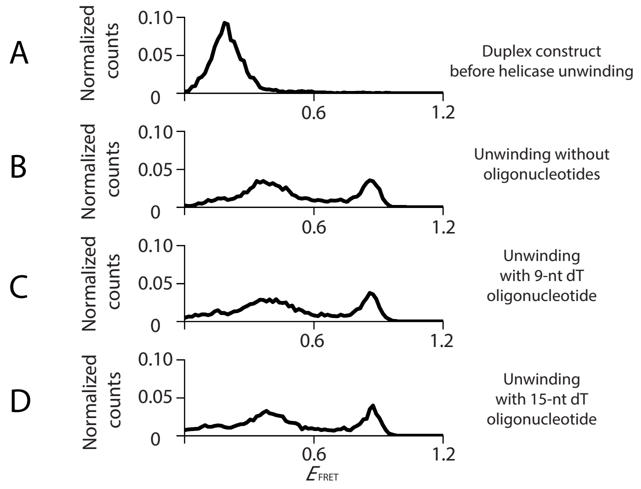


**Figure S10. The optimized unwinding condition for pseudo-functional studies during vectorial folding**.

In this condition, no observable loss of unwinding efficiency for addition of Rep-X along with free oligonucleotides. (**A**) Distribution of $E_{FRET}$ for heteroduplex before unwinding. (**B**) Distribution of $E_{FRET}$ after regular unwinding. (**C**) Distribution of $E_{FRET}$ for simultaneous addition of free dT9 oligonucleotides and extra Rep-X helicase. (**D**) Distribution of $E_{FRET}$ for simultaneous addition of free dT15 oligonucleotides and extra Rep-X helicase.

SUPPLEMENTARY TABLES

**Table S1. Sequence of mutations**

| Mutation | Sequence |
|---|---|
| Wild-typed | GAUACCUGUCACAACGGCU(U-Cy3)CCUGGCGUGACGAGGU GACCUCAGUGGAGCAAACCGCTGCCGTCGCTCCG |
| P1x_C26Z | GAUACCUGUCACAPCGGCU(U-Cy3)CCUGGCGUGACGAGGU GACCUCAGUGGAGCAAACCGCTGCCGTCGCTCCG |
| P1x_A14P | GAUACCUGUCACAACGGCU(U-Cy3)CCUGGZGUGACGAGGU GACCUCAGUGGAGCAAACCGCTGCCGTCGCTCCG |
| T_G16P | GAUACCUGUCACAACPGCU(U-Cy3)CCUGGCGUGACGAGGU GACCUCAGUGGAGCAAACCGCTGCCGTCGCTCCG |
| PK_C18A/G49U/C50U | GAUACCUGUCACAACGGAU(U-Cy3)CCUGGCGUGACGAGGU GACCUCAGUGGAUUAAACCGCTGCCGTCGCTCCG |
| P1x_A14C/C26U | GAUACCUGUCACACCGGCU(U-Cy3)CCUGGUGUGACGAGGU GACCUCAGUGGAGCAAACCGCTGCCGTCGCTCCG |
